# Financial strain and stressful social environment drive depressive symptoms, while *FKBP5* variant intensifies the effect, in African Americans living in Tallahassee

**DOI:** 10.1101/2020.07.16.206037

**Authors:** Kia Fuller, Clarence C. Gravlee, Chris McCarty, Miaisha M. Mitchell, Connie J. Mulligan

**Author notes:** Corresponding author (CJM).

## Abstract

The World Health Organization estimates that almost 300 million people suffer from depression worldwide. Depression is the most common mental health disorder and shows racial disparities in disease prevalence, age of onset, severity of symptoms, frequency of diagnosis, and treatment utilization across the United States. Since depression has both social and genetic risk factors, we propose a conceptual model wherein social stressors are primary risk factors for depression, but genetic variants increase or decrease individual susceptibility to the effects of the social stressors. Our research strategy incorporates both social and genetic data to investigate variation in symptoms of depression (CES-D scores). We collected data on financial strain (difficulty paying bills) and personal social networks (a model of an individual’s social environment), and we genotyped genetic variants in five genes involved in stress reactivity (*HTR1a, BDNF, GNB3, SLC6A4*, and *FKBP5*) in 135 African Americans residing in Tallahassee, Florida. We found that high financial strain and a high percentage of people in one’s social network who are a source of stress or worry were significantly associated with higher CES-D scores and explained more variation in CES-D scores than did genetic factors. Only one genetic variant (rs1360780 in *FKBP5*) was significantly associated with CES-D scores and only when the social stressors were included in the model. Interestingly, the effect of *FKPB5* appeared to be strongest in individuals with high financial strain such that participants with a T allele at rs1360780 in *FKBP5* and high financial strain had the highest mean CES-D scores in our study population. These results suggest that material disadvantage and a stressful social environment increases the risk of depression, but that individual-level genetic variation may increase susceptibility to the adverse health consequences of social stressors.

## Introduction

Depression is one of the most prevalent and most burdensome mental illnesses both in the United States (US) and globally. Approximately 17.3 million US adults and 264 million adults worldwide suffered from depressive disorders in 2017, which is more than any other mental disorder [1, 2]. Depression is characterized by loss of interest or pleasure in activities once enjoyed, feelings of guilt or low self-worth, disturbed sleep, lack of appetite, decreased ability to concentrate, and persistent feelings of sadness [3]. People with depression often suffer reduced quality of life. Furthermore, depression is the leading cause of disability in US adults among all mental illnesses, accounting for almost 400 million disability days each year [4]. Depression incurs a high economic burden, costing $210 billion in the US alone due to direct medical costs, suicide-related mortality costs, and indirect workplace costs [5].

Prevalence of depressive disorders, age of onset and severity of symptoms, as well as frequency of diagnosis and treatment show racial disparities across the US. African Americans have a lower prevalence of most forms of depression, such as major depression and persistent depressive disorder [4]. The exception is dysthymic disorder, a form of depression characterized by a low mood that lasts for at least two years with individuals feeling little to no joy. Lifetime prevalence of dysthymic disorder is 7.50% in African Americans and 5.70% in European Americans [6]. In addition, African Americans demonstrate a later age of onset but more severe symptoms of depression relative to European Americans [4], suggesting that diagnosis may be delayed in African Americans. Delays in diagnosis are likely due to multiple factors, including disparities in access to quality care [7], discrimination in healthcare settings, and a tendency not to seek care [8] because of long-term issues of trust as well as histories of stigmatization and exclusion of African Americans from US medicine [9].

Social conditions are fundamental causes of poor health, including depression [10-12]. In particular, social factors such as poverty, neighborhood conditions, racial discrimination, and cumulative material disadvantage over the life course are known contributors to racial and ethnic disparities in depression [13-15] and other mental illnesses. We interpret this evidence in light of conceptual models that highlight pathways of embodiment through which unequal material and social conditions become incorporated, literally, in biology [16, 17]. Here we integrate the focus on embodiment with Pluess and Belsky’s model of “vantage sensitivity.” Pluess and Belsky have proposed that genetic variants do not function solely as risk factors, but rather as environmental sensitivity variants whose carriers are more sensitive, or responsive, to both positive and negative exposures [18-20]. Thus, we propose that social and material conditions are primary risk factors for depression, but key genetic variants may increase the sensitivity of an individual to the effects of social stressors. In other words, poverty and social exclusion are bad for everyone, but the effects on mental health may be even worse for people with certain genetic variants, such as those that influence stress reactivity like the genes involved in the hypothalamic-pituitary-adrenal (HPA) axis. Studies of mental health that do not include both social and genetic data will fail to detect joint and independent effects of genetic and social factors, which may lead to undetected associations or conflicting results when compared across studies. Researchers have identified multiple genes associated with mental illness prevalence and severity, but the effect of single genes is typically small, e.g. 3% of variation in anxiety-related personality traits is attributable to the *SLC6A4* gene [21], leading to the possibility that larger effects between genetic and social factors have gone undetected.

Socioeconomic status (SES) and other measures of economic disadvantage have been established as significant risk factors for depressive disorders. The prevalence and persistence of depression are higher in individuals of economic disadvantage whether measured as poverty, SES, education, or income [10, 22-24]. The effects of economic disadvantage appear to be cumulative with those experiencing long term financial hardship being at the greatest risk of depression [23]. Moreover, prior depression has been associated with subsequent downward social mobility [24], demonstrating the persistent cycle of economic disadvantage and depression. Zimmerman and Katon [25] looked more in-depth at the relationship between economic disadvantage and depression and found that a measure of financial strain based on the ratio of debts-to-assets and employment status were both causally related to depression but income was not.

Depression is also influenced by an individual’s social environment, which can be investigated through social networks that model the social relationships in a person’s life. Ego-centered, or personal, social networks are constructed by asking study participants to list members of their social network (called ‘alters’) and report characteristics of each alter (such as age, sex, relationship to the study participant) and the likelihood that each alter knows every other alter. The characteristics of the network (e.g., average age of alters, number of women, number of family members, etc) can then be used to investigate outcomes of interest. Social networks have been shown to influence a range of mental and physical health outcomes, including happiness, obesity and blood pressure [26-28]. Rosenquist et al. [29] found that high and low scores on the Center for Epidemiological Studies Depression (CES-D) Scale were highly correlated with network members’ scores and that female friends were particularly influential. The ability of social networks to influence health through a type of social contagion is an understudied risk factor for mental health disorders.

Several genes (including the ones tested in this study: *HTR1a, BDNF, GNB3, SLC6A4*, and *FKBP5*) have been associated with depressive disorders although the results have not always been consistent across studies. *HTR1a* codes for the 5-hydrosytryptamine receptor 1a that is used in serotonin reuptake. The minor G allele of the rs6295 single-nucleotide polymorphism (SNP) in *HTR1A* has been associated with increased major depression and suicide in a linkage analysis of Utah pedigrees and a study of European American males [30, 31], but not in a study of males and females from Barcelona [32]. *BDNF* encodes the brain-derived neurotrophic factor (BDNF), a protein necessary for neuron development and survival in the central nervous system [33].

*BDNF* is thought to affect depression since BDNF supports the survival and protection of neurons that are necessary for hippocampal function and mood regulation [34], and there is suggestive evidence of a genetic effect on major depression [35]. *GNB3*, the guanine nucleotide binding protein β polypeptide 3 gene, produces a G protein messenger and is thought to aid in regulating dopaminergic activity. Multiple *GNB3* genetic variants have been found to associate with major depression in a Chinese population in combination with exposure to negative life events [36] and *GNB3* has been suggested as a possible target for antidepressant medication [37]. *SLC6A4* encodes the serotonin transporter protein, which is important in the inactivation and recycling of serotonin, which in turn helps maintain the release of serotonin and regulation of mood [38]. Variants in *SLC6A4* have been shown to predict depressive symptoms in African American women through an interaction with neighborhood crime [39] although another study of African and European Americans found no significant association between tested *SLC6A4* variants and major depression [40]. Finally, *FKBP5* encodes FK506 binding protein 5, which influences the HPA axis and reactivity of the HPA axis has been linked to mood disorders like depression [41]. Two SNPs (rs1360780 and rs3800373) have been associated with major depression in a meta-analysis [42] although reports of their effect on antidepressant medication response are mixed [43]. These SNPs have been primarily studied in European American populations, so study in African Americans is necessary to better understand the role of genetic variation in depressive disorders.

What remains unknown is how these factors—SES, social networks, and genetic variation—interact in the risk for poor mental health. In this study, we use a conceptual model for depression that explicitly integrates social and genetic risk factors for depression. We use this model to test the premise that financial strain and a stressful social environment increase the risk of depression, but that key genetic variants may influence susceptibility to these adverse social stressors. To accomplish this, we collected data on depressive symptoms (CES-D score), financial strain, and personal social networks, and we genotyped genetic variants in *HTR1a, BDNF, GNB3, SLC6A4*, and *FKBP5*. We used multiple linear regression and model building to identify the optimal model to explain variation in the output phenotype of CES-D score. The overarching hypothesis was that high levels of financial strain and a high percentage of stressful network members, in combination with genetic risk alleles, would be present in individuals with a higher number of depressive symptoms, as indicated by higher CES-D scores.

## Materials and Methods

### Ethics statement and data availability

The study protocol and informed consent procedure were approved by the University of Florida’s Institutional Review Boards (IRB-01 #364–2008 and IRB-02 #2007-U-469) and by the Health Equity Alliance of Tallahassee Steering Committee. Participants were asked to consent to give a saliva sample for genotyping purposes, to participate in the 2-3 hour sociocultural interview, to participate in the social network interview, and for their data to be used in other health-related studies besides blood pressure and cardiovascular disease. Participants were able to consent separately for each of these aspects of the study and only participants who consented to all four aspects were included in the study. Copies of the signed informed consents forms were scanned and uploaded to a secure online site accessible only by CCG and CJM. All data analyzed in this study are available at https://data.mendeley.com/datasets/3f46kg3m55/1.

### Study population

Data for this study come from the HEAT Heart Health study, a community-based participatory research (CBPR) project of the Health Equity Alliance of Tallahassee (HEAT). Study participants were selected from a multistage probability sample of African Americans living in Tallahassee, Florida, as described in Fuller et al. [28]. Cluster analysis of neighborhood level markers of socioeconomic status and racial composition were used to group census blocks.

Postal addresses were then selected randomly from groups of census blocks. Participants were able to participate if they were at least eighteen years old. The initial sample size of this study was n=185. Twenty participants did not consent to give saliva samples and 19 participants did not agree to participate in the second interview, which collected the social network data. After accounting for DNA samples that did not type for all genetic variants and removing participants with incomplete data, the final sample size for this study was n=135 participants.

### Measuring symptoms of depression and financial strain

Symptoms of depression in participants were measured using the Center for Epidemiologic Studies Depression (CES-D) scale. The CES-D is a validated and widely used self-administered tool that evaluates symptoms of depression in an individual [44]. The CES-D is a 20-item questionnaire with a maximum score of 60. The American Psychological Association (APA) classifies a score ≥ 16, the standard cutoff point, as an indication of the potential for clinical depression [45]. The potential responses to the CES-D questions are “rarely or none of the time (less than 1 day), “some or a little of the time (1-2 days)”, “occasionally or a moderate amount of time (3-4 days)”, and “most or all of the time (5-7 days). In our study, the CES-D form was modified, by separating one of the existing responses (“rarely or none of the time”) into two responses (“rarely” and “none of the time”), in order to capture a wider range of responses, which then increased the maximum possible score from 60 to 80. When using the validated cutoff of ≥ 16 for risk of clinical depression, the responses were translated back to their original form, i.e. “rarely” and “none of the time” were grouped into the original response of “rarely or none of the time”.

In keeping with CBPR principles, interviews were conducted by members of the Tallahassee community who were trained as interviewers. CES-D data were collected during the first of two interviews. The first interview lasted approximately 2-3 hours and collected a wealth of sociocultural data including: age, sex, financial strain, neighborhood ecology, and social stressors. In this study, we focused on financial strain as a measure of financial stress and socioeconomic status, which are factors that have been associated with depression in other studies [23]. Financial strain was dichotomized for analyses, i.e. “very difficult to pay bills” plus “somewhat difficult” versus “not very difficult” plus “not at all difficult”.

### Social networks

Social network data were collected during the second interview using the EgoNet program [46], which also constructed and visualized the social networks. All participants were required to name 30 alters, or members of their network, using the following prompt: *“Name 30 people that you know by sight or by name, whom you could contact today if you needed to”*. Participants were then asked to provide information on each of their 30 alters, including age, sex, their relationship with the person, and how often they feel the person causes them stress or worry. We also elicited information about the structure of the network by asking participants to estimate the ties among members of their network.

In this study, we focus on responses to the question, “Does [name of alter] ever cause you any stress or worry?” Response options included “never,” “hardly ever,” “sometimes,” “often,” and “always.” To quantify the level of stress in one’s proximate social environment, we calculated the percentage of alters (out of 30) in each participant’s network who were “often” or “always” a source or stress or worry. For shorthand, we refer to this measure as social network stress.

### Genotyping

Saliva samples were collected and stored using Oragene saliva collection kits (DNA Genotek, Ontario, Canada). Extraction of DNA was performed according to the manufacturer’s protocol. Samples were genotyped for the following SNPs: rs6295 (*HTR1a*), rs4922793 (*BDNF*), rs5443 (*GNB3*), rs140701 (*SLC6A4*), and rs1360780 (*FKBP5*). Three SNPs (rs4922793, rs140701, and rs1360780) were genotyped using a custom Affymetrix Axiom Array (Affymetrix, Santa Clara, CA) designed by Laurel N. Pearson. Two SNPs (rs6295 [assay ID: C_11904666_10] and rs5443 [assay ID: C_2184734_10]) were genotyped using TaqMan® Genotyping Assays following manufacturer protocols (Thermo Fisher Scientific Inc, Waltham, MA). Genotypes for all SNPs were determined to be in Hardy-Weinberg equilibrium through testing using the Hardy-Weinberg equilibrium calculator tool [47].

### Statistical analysis

#### Model building

Models were built to determine which factors contributed to variation in the output phenotype of depressive symptoms as measured by CES-D score. Multiple linear regression and Akaike Information Criterion (AIC) measures were used to develop models to describe the variation in CES-D scores observed in the sample population. A stepwise AIC provides a measure of relative model quality by sequentially adding and removing variables, generating new models, and comparing each new model to the original. Four models were generated to explain variance in CES-D score. The first model included only the standard covariates age and sex. The second, called the genetic model, included the standard covariates plus data on the five tested SNPs. The third model, called the social model, included the standard covariates plus financial strain (difficulty paying bills) and social network stress. The fourth model included the standard covariates and both the genetic and social data described above. The False Discovery Rate (FDR) method was used to correct for multiple comparisons in our hypothesis testing; after adjusting by FDR, a q-value ≤ 0.1 was used to indicate statistical significance.

#### Identification of the optimal CES-D model

Adjusted R^2^ values and AIC scores were used to evaluate the fit of the models. Lower AIC scores and higher adjusted R^2^ values indicate a model that is a relatively better fit for the data than a model with higher AIC scores and lower adjusted R^2^ values. Variables retained in the optimal model were included in the multiple linear regression table even if they were not found to be statistically significant since they improved AIC scores and R^2^ values. Analysis of Variance (ANOVA) was used to test the optimal model for statistically significant improvement against the genetic and social models.

## Results

### Study demographics

Demographic information on study participants is reported in Table 1. The mean age of all study participants was 41.3 years old. Women represented approximately 67% of the study population. The mean CES-D score for all study participants was 12.7. Average financial strain was approximately 1.5, indicating there were relatively equal numbers of participants who said that bill paying was “Not very difficult” or “Not at all difficult” (coded as 2) and those who said that bill paying was “Very difficult” or “Somewhat difficult” (coded as 1). As a measure of social network stress, study participants said that 5-6% of their social network members caused them stress or worry “often” or “always”. There was no significant difference in mean CES-D scores between males and females (14.5 vs 11.8, Pr(>F) =0.09). The genotype counts for the tested SNPs (rs140701, rs1360780, rs4922793, rs5443, and rs6295) are detailed in Table 2.

**Table 1.** Mean characteristics of study participants.

| Characteristics | Male | Female | Total Sample |
| --- | --- | --- | --- |
| N | 45 | 90 | 135 |
| Age, years (SD) | 40.2 (12.2) | 41.8 (11.7) | 41.3 (11.8) |
| **CES-D score (SD)** | 14.5 (9.34) | 11.84(8.20) | 12.7 (8.7) |
| **Financial Strain** | 1.51 (0.50) | 1.54 (0.50) | 1.53 (0.50) |
| **Social network stress** | 6% (8%) | 5% (8%) | 5% (7%) |
| **(percent of alters)** |  |  |  |

**Table 2.** Genotype counts for rs140701, rs1360780, rs4922793, rs5443, rs6295.

| **Gene/SNP** | **Genotype** | **Number of individuals** |
| --- | --- | --- |
| ***SLC6A4/rs140701*** | AA | 13 |
|  | AG | 65 |
|  | GG | 57 |
| ***FKBP5/rs1360780*** | CC | 26 |
|  | CT | 60 |
|  | TT | 49 |
| ***BDNF/rs4922793*** | AA | 120 |
|  | AG | 15 |
|  | GG | 0 |
| ***GNB3/rs5443*** | TT | 73 |
|  | TC | 54 |
|  | CC | 7 |
| ***HTR1AI/rs6295*** | CC | 32 |
|  | CG | 58 |
|  | GG | 44 |

### Multiple linear regression

Four models were tested in order to identify the optimal model that explained the most variation in CES-D score (Table 3). The model with the highest adjusted R^2^ and lowest AIC score was chosen as the optimal model, which was the fourth model that contained both genetic and social data (Table 3). The optimal model accounted for 19% of variation (adjusted R^2^) in CES-D scores in comparison to the genetic model and social models, which explained 2% and 16% of CES-D score variation, respectively. The variables included in the optimal model were financial strain, social network stress, and SNPs in *SLC6A4* and *FKBP5*. Of the variables included in the optimal model, all except the *SLC6A4* SNP were significantly associated with CES-D score (Table 3).

**Table 3.** Multiple linear regression models including genetic and social variables to predict CES-D scores*.

|  | **Standard**<br>**Covariate Model** |  |  | **Genetic Model** |  |  | **Social Model** |  |  | **Optimal Model** |  |  |
| --- | --- | --- | --- | --- | --- | --- | --- | --- | --- | --- | --- | --- |
| Variable | Co-<br>effic<br>ient | Std.<br>Error | Q-value | Co-<br>effici<br>ent | Std.<br>Error | Q-valu<br>e | Co-<br>effici<br>ent | Std.<br>Erro<br>r | Q-value | Co-<br>effici<br>ent | Std.<br>Error | Q-value |
| Age | 0.02 | 0.06 | 0.71 | 0.03 | 0.06 | 0.78 | -0.01 | 0.06 | 0.92 | -0.01 | 0.06 | 0.88 |
| Sex<br>(male =1,<br>female =<br>2) | 1.36 |  | 0.20 | -3.34 | 1.69 | 0.18 | -2.04 | 1.47 | 0.22 | -2.52 | 1.45 | 0.10 |
| *HTR1A*<br>rs6295 |  |  |  | 0.52 | 0.99 | 0.78 |  |  |  |  |  |  |
| *GNB3*<br>rs5443 |  |  |  | -0.10 | 0.92 | 0.92 |  |  |  |  |  |  |
| *BDNF*<br>rs4922793 |  |  |  | 1.45 | 2.36 | 0.78 |  |  |  |  |  |  |
| *SLC6A4*<br>rs140701 |  |  |  | 2.24 | 1.16 | 0.18 |  |  |  | 1.87 | 1.06 | 0.101 |
| *FKBP5*<br>rs1360780 |  |  |  | 1.87 | 1.05 | 0.18 |  |  |  | **1.98** | **0.94** | **0.08** |
| Percent of<br>alters<br>rated as |  |  |  |  |  |  | **35.05** | **8.78** | **0.0004** | **34.81** | **8.63** | **0.001** |
| highly stressful |  |  |  |  |  |  |  |  |  |  |  |  |
| Financial strain |  |  |  |  |  |  | **4.19** | **1.71** | **0.03** | **4.18** | **1.72** | **0.05** |
| Adjusted R<sup>2</sup> | 0.02 |  |  | 0.02 |  |  | 0.16 |  |  | 0.19 |  |  |
| AIC Value | 585 |  |  | 588 |  |  | 563 |  |  | 559 |  |  |

\*Statistically significant findings are shown in bold font, q-value $\leq 0.10$

In the optimal model, high financial strain (“very difficult to pay bills” plus “somewhat difficult to pay bills”) was associated with higher CES-D scores (q-value = 0.05) and participants who reported a larger percentage of alters in their social network who were “often” or “always” a source or stress or worry also had higher CES-D scores than their peers (q-value ≤ 0.001). Both of these variables were significantly associated with CES-D score in the social model and the optimal model (Table 3). Social network stress was found to have the largest beta value and the smallest p-value, and therefore explained the most variation in depressive symptoms in our sample.

Of the genetic data, only one SNP (rs1360780 in *FKBP5*) was significantly associated with depressive symptoms and only when included in a model with social data. Furthermore, the *FKBP5* SNP had the smallest beta value (i.e. explained the least variation) of the three risk factors significantly associated with CES-D score. To better understand why the *FKBP5* SNP emerged as significant only when included in a model with social data, we tested for statistical interactions between *FKBP5* and financial strain or social network stress, but neither of these interactions was significant. We then plotted the effect of the *FKBP5* SNP on both social stressors and found an interesting pattern with financial strain. As shown in Fig 1, participants with high financial strain showed higher mean CES-D scores than individuals with low financial strain. Furthermore, individuals who were heterozygous or homozygous for the T allele at rs1360780 in *FKBP5* and reported high levels of financial strain had the highest mean CES-D scores in our study population. Specifically, individuals with a T allele and high financial strain had higher larger increases in mean CES-D scores than individuals with a T allele and low financial strain.

**Fig 1:**
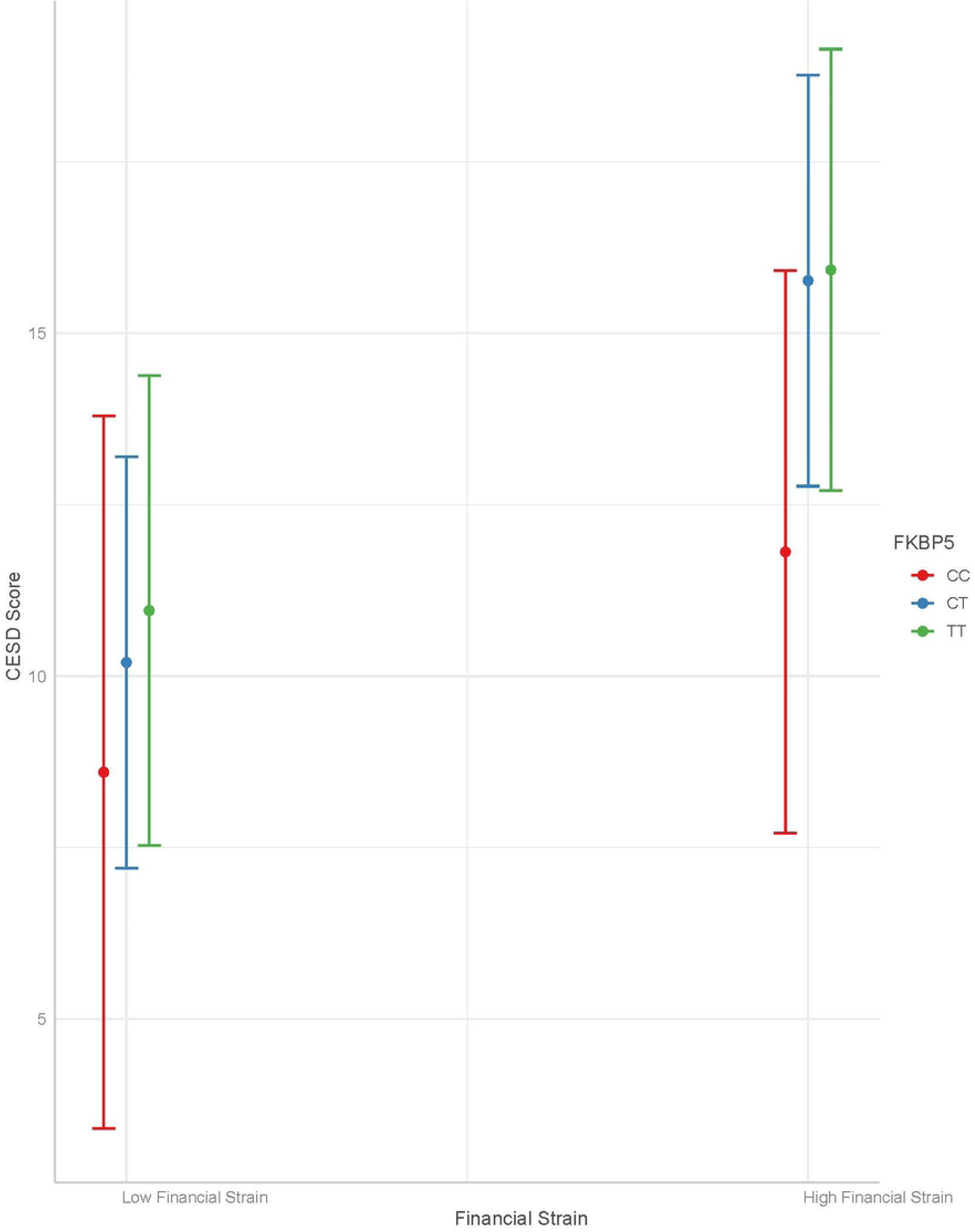
Interaction plot of *FKBP* rs1360790 genotype and financial strain with mean CES-D score. ‘Low Financial Strain’ represents participants who said bill paying was “Not very difficult” or “Not at all difficult” and ‘High Financial Strain’ represents participants who said bill paying was “Very difficult” or “Somewhat difficult”. Colored lines represent the three possible *FKBP5* genotypes. Dots represent the mean CES-D score and bars represent the variance in CES-D score.

## Discussion

Overall, we observed that social risk factors involving stress due to financial strain and a stressful social environment were significantly associated with symptoms of depression, as measured by CES-D scores in our sample of African Americans in Tallahassee, FL (Table 3). Previous studies have linked depressive symptoms to social stressors related to poverty and social network characteristics [23-25, 29]. We measured these stressors as difficulty paying bills (financial strain) and having a high percentage of people in one’s social network who cause stress or worry (social network stress). Social network stress accounted for more variation in depressive symptoms than did any other variable in our study. We simultaneously investigated the associations between depressive symptoms and five genetic variants, or SNPs. Only one of the SNPs, rs1360780 at *FKBP5*, was significantly associated with CES-D scores, and only when financial strain and social network stress were included in the model. In the model excluding social data, none of the genetic variants was associated with depressive symptoms, and in the optimal model including social data, the *FKBP5* SNP explained the smallest amount of variation in CES-D scores.

In order to better understand why the *FKBP5* SNP was only significant in the presence of the social stressors, we noted an interesting interplay between financial strain and the *FKBP5* SNP where individuals with the T allele had larger increases in CES-D scores associated with high financial strain than did individuals who were homozygous for the C allele (Figure 1). Our results are consistent with Pluess and Belsky’s model of environmental sensitivity that posits carriers of sensitivity variants are more sensitive to both positive and negative environmental exposures [18, 20, 48]. In our study, carriers of the T allele at rs1360780 in *FKBP5* appear to be more sensitive to the effects of financial strain on the risk of depressive symptoms than individuals who do not carry the T allele.

Poverty is generally associated with increased prevalence of poor physical and mental health, although the best measure of economic disadvantage and the mechanism by which such disadvantage acts are not clear. In general, income appears to be less impactful than the ability to pay bills, as is captured in Zimmerman and Katon’s measure of financial strain that is used in this study and has been causally related to symptoms of depression [25]. Specifically, we found that participants who reported feeling that it was “very difficult” or “somewhat difficult” to pay bills every month had significantly higher CES-D scores than those who responded otherwise. The mechanism by which material disadvantage impacts mental health is more difficult to determine. The fact that 10-20% of the US population lives at or below the poverty level [49], but not all experience depression, indicates, as we would expect for a complex phenotype, that factors other than poverty are involved in developing depression. Pluess [19] proposed a theoretical framework in which sensitivity to environmental stressors, like poverty, is influenced by specific genetic variants, and that genetic factors can help predict individual differences in environmental sensitivity to both positive and adverse conditions. Based on our results, presence of a T allele at rs1360780 in *FKBP5* and high levels of financial strain may predispose individuals living in poverty to be at increased risk for depression.

In order to better understand how *FKBP5* variation may influence sensitivity to financial strain, it is important to understand how the *FKBP5* gene product functions. FKBP5 regulates the sensitivity of the glucocorticoid receptor (GR) by inhibiting GR binding affinity in response to stress [50]. FKBP5 and GR are involved in a negative feedback loop in which FKBP5 inhibits GR sensitivity and GR controls *FKBP5* transcription [51]. Chronic stress leads to elevated glucocorticoid levels and specific *FKBP5* variants lead to increased *FKBP5* expression, which together can disrupt the delicate balance between GR and FKBP5 levels. Such an imbalance can result in impaired cognitive processing and increase the risk of stress-related mental health disorders, like depression and PTSD, as has been found for the *FKBP5* rs1360780 variant [52, 53]. With respect to our results, the over-expression associated with the T allele of rs1360780 in *FKBP5* may create an imbalance between FKBP5 and GR levels that is further exacerbated by the social stress caused by financial strain.

Binder et al. (2004) first investigated the effect of *FKBP5* variants on depression and found no association with the disease, but did identify a significant association between *FKBP5* variants and recurrence of depressive episodes and rapid response to antidepressants [53]. More recently, Ising et al. (2019) [54] found that the increased levels of *FKBP5* RNA and protein found in depressed patients were reduced with antidepressant use and this effect was most pronounced in carriers of the T allele at rs1360780, suggesting that changes in FKBP5 levels are causative in terms of development and treatment of depression and that the T allele is most responsive to the environment (in this case, antidepressant use). Recent studies have found interesting interactions between the T allele and early life stress or adversity similar to our finding of increased CES-D scores in T allele carriers who experience high financial strain. Specifically, a recent meta-analyses found that carriers of the T allele at rs1360780 who were exposed to early life stress were at higher risk for developing depression [55]. Zimmerman et al. found no main genetic effect of rs1360780, but identified an interaction between the T allele and exposure to traumatic events that was significantly associated with occurrence of depressive episodes [56]. Our results suggest that material disadvantage, such as financial strain, should also be considered a type of stress that can interact with the *FKBP5* variant to increase risk of depression.

Our study also revealed that characteristics of one’s social network associated with the number of depressive symptoms. Specifically, having a higher percentage of network members who are a cause of stress or worry was significantly associated with higher CES-D scores (Table 3). This measure of social network stress was the largest contributor to variance accounted for in our model. Social networks are being used more frequently to quantify aspects of an individual’s social environment. Variation in network composition and structure have been linked to a range of outcomes including happiness, obesity, and blood pressure [26-28]. Researchers typically invoke social networks as a source of social support, but personal networks can also be a source of strain. In a previous study with the same population of African Americans, we found that systolic and diastolic blood pressure were significantly elevated in individuals with networks dominated by family members [28]. In the current study, we show that depression is associated with the percentage of the network comprised of people who cause participants stress or worry. We interpret these findings, together, as evidence of the specifically *social* nature of social stressors. The burdens people carry extend beyond their own experiences to the worries of others [57]. We cannot tell from our data what the specific worries are—concern about sheltering children from racism, caring for loved ones who are sick, recovering from an abusive relationship, assisting family members experiencing financial strain of their own [58] – but we suggest that high levels of network stress reflect one’s position in systems of inequality. That is, social networks may connect macro-scale social forces like poverty and racism to individual-level health and well-being. Future research should seek to clarify which dimensions of personal networks constitute resilience resources and which add to the cumulative burden of social stressors that increases the risk for poor health outcomes like depression [59, 60].

There are several limitations to note in our study. We were able to detect important associations between financial stress, social network stress, and the rs1360780 variant at *FKBP5* with symptoms of depression; however, we were limited by a relatively small sample population that could have impacted our ability to detect more complex associations or interactions. Furthermore, we lacked data on participant history of being diagnosed with or treated for depression, which meant that we could not account for history of depression in our analyses. Future studies should test larger sample sizes for the impact of social and material risk factors on depression and the role of genetic variants to modify susceptibility to such risk factors. Finally, in order to investigate the interplay between specific social and genetic risk factors, we tested a model that consisted of two social stressors and five genetic variants so we cannot comment on the contribution of other risk factors for depressions.

## Conclusions

We test a conceptual model wherein social stressors are primary risk factors for depression, while genetic variants increase or decrease susceptibility to the effects of social stressors. We demonstrate that high financial strain and a high percentage of stressful individuals in one’s social network significantly associated with higher CES-D scores and explained most of the variation in CES-D scores in our study population. Of the five tested genetic variants, only one (rs1360780 in *FKBP5*) was significantly associated with CES-D scores and only when social stressors were included in the model. Specifically, participants with high financial strain and a T allele at rs1360780 in *FKBP5* had the highest mean CES-D scores in our study population. These results suggest that material disadvantage (measured here as financial strain) increases the risk of depression, but that individual-level genetic variation may increase susceptibility to the adverse health consequences of such material disadvantage. A corollary of this finding is that genetic influences on the risk of depression, which appear to be small at the population level, cannot be understood without attention to the social and material conditions in which people live.

## Acknowledgements

We thank our study participants for their contributions. We thank Malika Macey, Samantha McCrane, Jess Ross, and Olivia Trumble for their assistance with genotyping the genetic variants. The research is a project of the Health Equity Alliance of Tallahassee (HEAT), a community-academic partnership for action-oriented research on social inequalities in health (http://healthequityalliance.org/). We thank HEAT Steering Committee members Mr. James Bellamy, Dr. Qasimah Boston, Dr. Edward Holifield, and Dr. Cynthia Seaborn for their insight and guidance. The study was supported by NSF grants BCS 0820687 and BCS 0724032.

